# Human IgE producing B cells have a unique transcriptional program and generate high affinity, allergen-specific antibodies

**DOI:** 10.1101/327866

**Authors:** Derek Croote, Spyros Darmanis, Kari C. Nadeau, Stephen R. Quake

## Abstract

IgE antibodies provide defense against helminth infections, but can also cause life-threatening allergic reactions. Despite their importance to human health, these antibodies and the cells that produce them remain enigmatic due to their scarcity in humans; much of our knowledge of their properties is derived from model organisms. Here we describe the isolation of IgE producing B cells from the blood of individuals with food allergies, followed by a detailed study of their properties by single cell RNA sequencing (scRNA-seq). We discovered that IgE B cells are deficient in membrane immunoglobulin expression and that the IgE plasmablast state is more immature than that of other antibody producing cells. Through recombinant expression of monoclonal antibodies derived from single cells, we identified IgE antibodies which had unexpected cross-reactive specificity for major peanut allergens Ara h 2 and Ara h 3; not only are these among the highest affinity native human antibodies discovered to date, they represent a surprising example of convergent evolution in unrelated individuals who independently evolved nearly identical antibodies. Finally, we discovered that splicing within B cells of all isotypes reveals polarized germline transcription of the IgE, but not IgG4, isotype as well as several examples of biallelic expression of germline transcripts. Our results offer insights into IgE B cell transcriptomics, clonality and regulation, provide a striking example of adaptive immune convergence, and offer an approach for accelerating mechanistic disease understanding by characterizing a rare B cell population underlying IgE-mediated disease at single cell resolution.

## Introduction

The IgE antibody class is the least abundant of all isotypes in humans and plays an important role in host defense against parasitic worm infections (1), but it can also become misdirected towards otherwise harmless antigens. Food allergies are one example of this misdirection, where the recognition of allergenic food proteins by IgE antibodies can lead to symptoms ranging from urticaria to potentially fatal anaphylaxis. Despite this central role in immunity and allergic disease, human IgE antibodies remain poorly characterized due to their scarcity (2). Bulk epitope mapping experiments have revealed that IgE antibodies are polyclonal and epitopes are heterogeneous (3); however, individuals with the same allergy tend to recognize a core set of one or a few allergenic proteins (4). Recent studies applying bulk fluorescence activated cell sorting (FACS) immunophenotyping (5, 6) and immune repertoire deep sequencing (7) have inferred IgE B cell characteristics and origins, while studies performing peanut allergen specific single cell sorting (8, 9) have described clonal families to which IgE antibodies belong. However, none have successfully isolated single IgE producing cells or the paired heavy and light chain sequences that comprise individual IgE antibodies, leaving unanswered questions as to the functional properties of such antibodies, the transcriptional programs of these cells, and the degree to which any of these features are shared across individuals. Similarly, there is a lack of knowledge, but growing interest, surrounding the IgG4 isotype due to its potential to compete with IgE for allergen and thereby contribute to the reduced clinical allergen reactivity that accompanies immunotherapy and early allergen exposure (10). Here we report the first successful isolation and transcriptomic characterization of single IgE and IgG4 producing B cells from humans. We combined single cell RNA sequencing (scRNA-seq) with functional antibody assays to elucidate mechanisms underlying the regulation of IgE and to discover high affinity peanut-specific antibodies.

### Characterization of single B cells from peripheral blood

We performed scRNA-seq on B cells isolated from the peripheral blood of food allergic individuals, which enabled us to characterize each cell’s gene expression, splice variants, and heavy and light chain antibody sequences (Fig. 1A). Fresh peripheral blood from six peanut allergic individuals was first separated into plasma and cellular fractions; plasma was stored and later used for allergen-specific IgE concentration measurements (fig. S1), while the cellular fraction was enriched for B cells prior to FACS (see Materials and Methods). CD19+ B cells of all isotypes were sorted exclusively based on immunoglobulin surface expression, but with an emphasis on maximizing IgE B cell capture (fig. S2). Because cellular isotype identity was determined post-hoc from scRNA-seq, we were able to sacrifice specificity and capture cells with high sensitivity. This approach makes the prospect of IgE B cell capture accessible for many laboratories without stringent requirements on FACS gate purity or the need for complex, many-color gating schemes.

**Fig 1.**
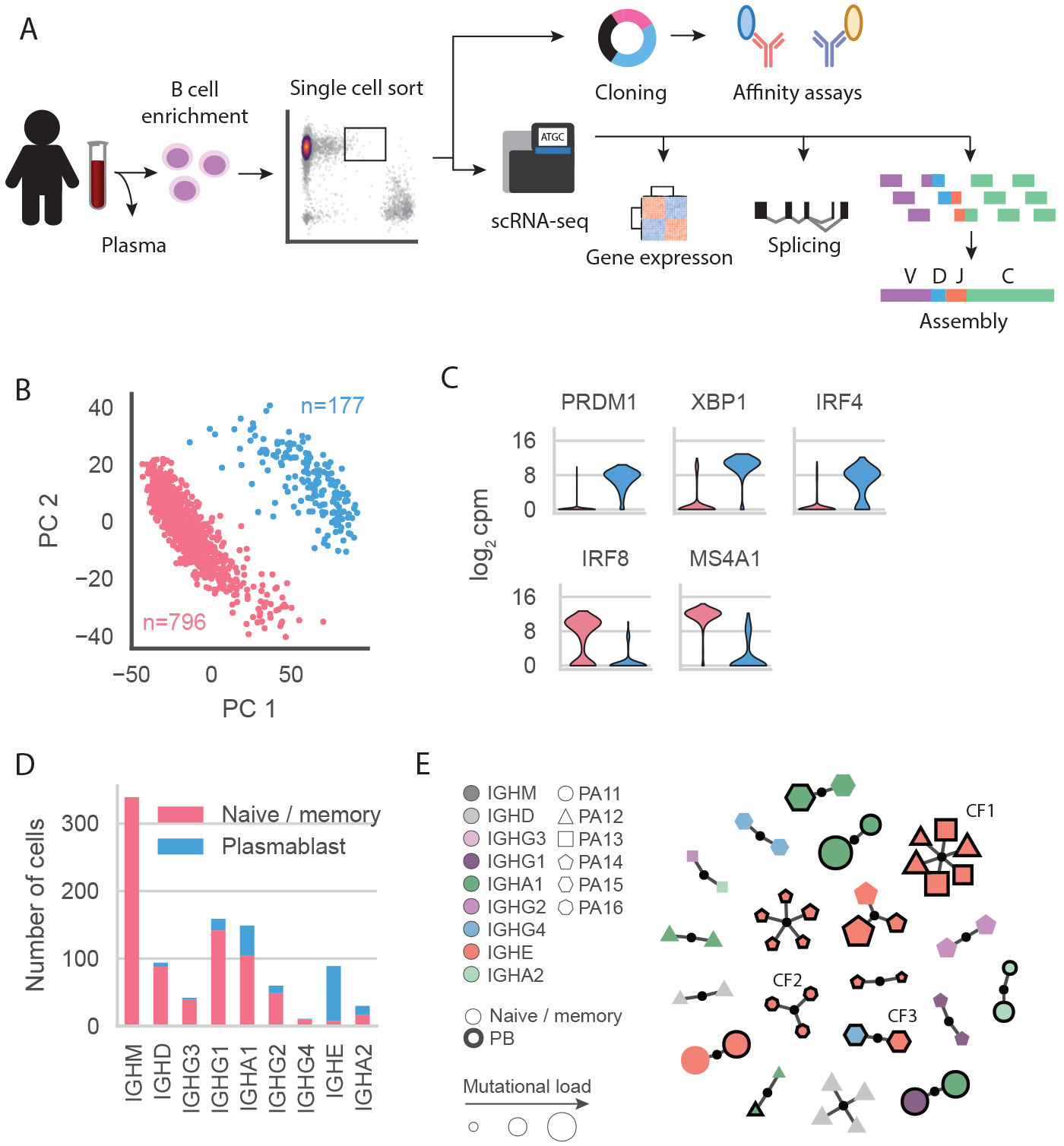
Characterization of single B cells isolated from fresh peripheral blood of allergic individuals. (A) Study overview. (B) Principal component analysis separates naïve / memory (pink) and plasmablast (PB, blue) B cell subsets identified by expression of established transcription factors and marker genes shown in (C). (D) Number of cells belonging to each subtype in (B) by isotype. (E) Isotype, B cell subtype, patient of origin, and mutational frequency of each cell that belongs to a clonal family (CF) with multiple members. CFs referred to in the text are labeled.

Single cells were sorted into 96 well plates, processed using a modified version of the Smart-seq2 protocol (11), and sequenced on an Illumina NextSeq 500 with 2×150 bp reads to an average depth of 1-2 million reads per cell (fig. S3). Sequencing reads were independently aligned and assembled to produce a gene expression count table and reconstruct antibody heavy and light chains, respectively (see Materials and Methods). Using STAR (12) for alignment also facilitated the assessment of splicing within single cells. Cells were stringently filtered to remove those of low quality, putative basophils, and those lacking a single productive heavy and light chain, yielding a total of 973 cells for further analysis. The isotype identity of each cell was determined by its productive heavy chain assembly, which avoids misclassification of isotype based on FACS immunoglobulin surface staining (fig. S2B), a problem which is especially pervasive for IgE B cells due to CD23, the “low-affinity” IgE receptor that captures IgE on the surface of non-IgE B cells (6).

Principal component analysis of normalized gene expression following batch effect correction (fig. S3 and Materials and Methods) separated cells into two distinct clusters identifiable as plasmablasts (PBs) and naïve / memory B cells (Fig. 1B-C). PBs expressed the triad of transcription factors BLIMP1 (PRDM1), XBP1, and IRF4 that drive plasma cell differentiation (13), as well as genes associated with antibody secretion (fig. S4), while naïve and memory cells expressed the canonical mature B cell surface marker CD20 (MS4A1), as well as transcription factor IRF8, which antagonizes the PB fate and instead promotes a germinal center response (14). Additional data corroborated this cell subtype assignment; PBs had greater FACS forward and side scatter in agreement with their larger size and increased granularity, PB cDNA concentrations were higher following preamplification, and PBs expressed more antibody heavy and light chain transcripts (fig. S4).

We assessed the distribution of isotypes within each B cell subtype and found that, in stark contrast to other isotypes, circulating IgE B cells overwhelmingly belonged to the PB subtype (Fig 1D, fig. S5A). This discovery is consistent with observations of preferential differentiation of IgE B cells into PBs in mice (15). Subtype proportions for other isotypes followed expectations: IgM B cells, which are primarily naïve, had the lowest PB percentage, while IgA B cells had the highest in accordance with their secretory role in maintaining mucosal homeostasis. Interestingly, we found that the number of circulating IgE B cells for each individual correlated with total plasma IgE levels (fig S1C); a similar phenomenon has been noted in atopic individuals and individuals with hyper-IgE syndrome (16).

By clustering antibodies into clonal families (CFs) we were able to observe elements of classical germinal center phenomena such as somatic hypermutation, class switching, and fate determination in our data. Using a standard immune repertoire sequencing approach (17), all antibody heavy chain sequences were first divided by V and J genes and were clustered if their amino acid CDR3 sequences shared at least 75% similarity. Only 49 heavy chains formed CFs with multiple members, although this was not surprising given the vast diversity of potential immunoglobulin gene rearrangements (fig. S5B). Within multi-member CFs, light chains were highly similar (fig. S5D), while overall, multi-member CFs were diverse (Fig. 1E); they contained between two and six sequences, had variable isotype membership, and had a comprehensive distribution of mutational frequency. CFs were specific to an individual, with the exception of one CF (CF1) that contained six heavily mutated IgE PBs: three each from individuals PA12 and PA13, as discussed in depth later. Four CFs illustrated the two possible B cell differentiation pathways in that they contained both PBs and memory B cells. Other CFs contained cells belonging to multiple isotypes, with one of particular interest (CF3), discussed later, that contained an IgE PB and an IgG4 PB. Interestingly, we found that in contrast to other isotypes, IgE and IgG4 were surprisingly clonal as over 20% of IgE and IgG4 cells belonged to such multi-member CFs (fig. S5C).

Across all individuals, the 89 IgE antibodies we found varied widely in gene usage and mutation frequency (Fig. 2A). They also varied in heavy and light chain CDR3 lengths (fig. S6A). There was moderate correlation between the mutation frequency of heavy and light chains within single cells (fig. S6B), with evidence of selection via an enrichment of replacement mutations relative to silent mutations in the heavy chain CDR1 and CDR2 that was absent in framework (FWR) regions. Light chains were similarly enriched for replacement mutations in the CDR1 and, to a lesser degree, FWR2 (fig. S6C). Compared to other isotypes, IgE B cells had a similar distribution of heavy chain mutation frequency, relative utilization of the lambda versus kappa light chains, and heavy chain V and J gene usage (fig. S6D-F).

**Fig 2.**
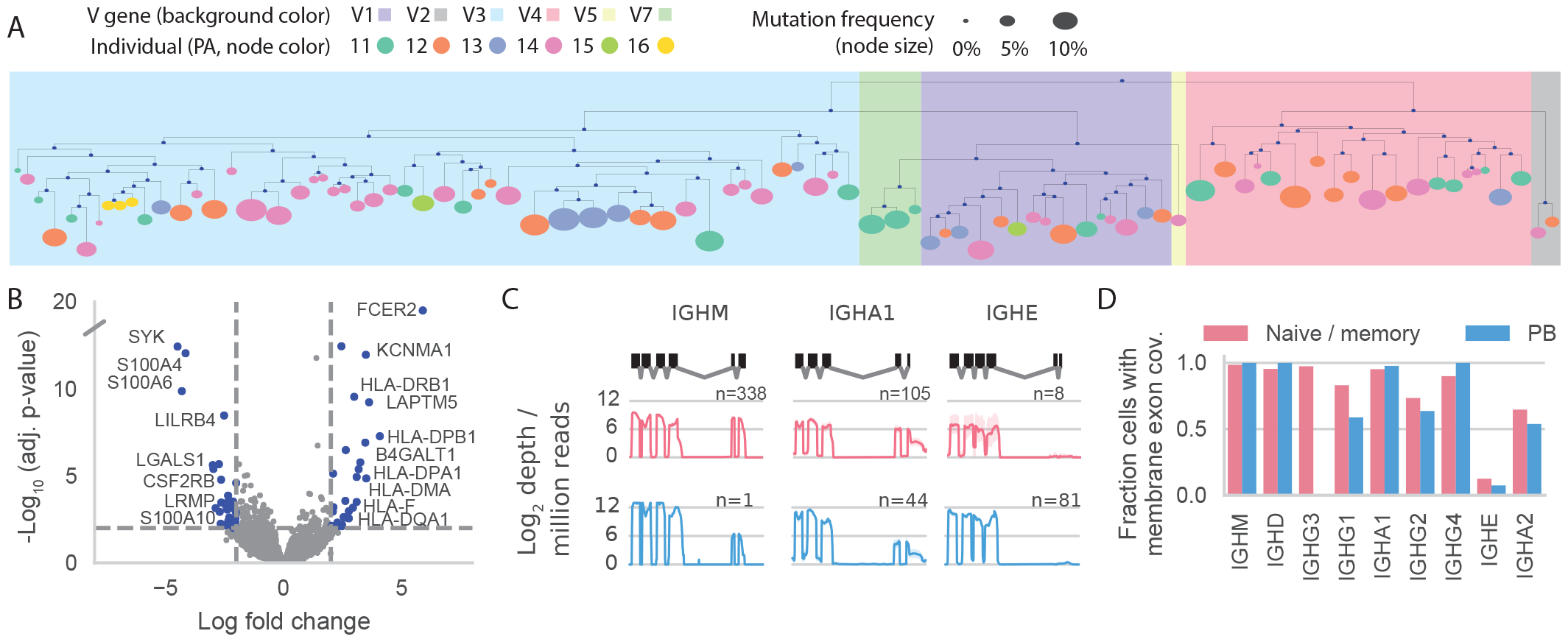
Characterization of 89 IgE antibodies and the single B cells that produce them. (A) Phylogenetic depiction of antibody heavy chains arranged by IGHV gene (background color), patient of origin (node color), and mutation frequency (node size). (B) Differential gene expression between IgE PBs and PBs of other isotypes. Positive log fold change indicates genes enriched in IgE PBs. (C) Heavy chain constant region coverage histograms for naïve / memory B cells (top) and PBs (bottom) for select isotypes. Mean normalized read depth and 95% confidence interval are indicated by solid lines and shaded area, respectively, for the number of cells (n) inscribed. Heavy chains are oriented in the 5’ to 3’ direction and membrane exons are the two most 3’ exons of each isotype. (D) Summary of (C), but depicting the fraction of cells of each isotype with any membrane exon coverage for both B cell subsets and all isotypes.

### IgE B cells possess a unique transcriptional program

To elucidate B cell intrinsic factors affecting PB activation, survival, and differentiation, we assessed genes differentially expressed between IgE PBs and PBs of other isotypes (Fig. 2B).

A host of MHC genes were robustly upregulated in IgE PBs, suggesting a more immature transcriptional program given the established loss of MHC-II during the maturation of PBs to plasma cells (18–20). FCER2 (CD23), the “low-affinity” IgE receptor was also highly upregulated, although its precise role within IgE PBs is unclear; autoinhibition of IgE production could result from membrane CD23-mediated co-ligation of membrane IgE (mIgE) and CD21 (21). Alternatively, IgE production could be upregulated by soluble CD23 (22), which is produced following cleavage by ADAM10 (23), a disintegrin and metalloproteinase domain-containing protein that we find is co-expressed in a subset of IgE PBs. LAPTM5, a negative regulator of B cell activation, BCR expression, and antibody production (24), was also upregulated, while CSF2RB, which encodes the common beta chain of the IL-3 and IL-5 receptors, was downregulated, potentially indicating weakened IL-3- and IL-5-mediated terminal differentiation capacity (25, 26). Additional downregulated genes included galectin 1 (LGALS1), which supports plasma cell survival (27) and the S100 proteins S100A4, S100A6, and S100A10, which may indicate reduced proliferative and survival signaling (28, 29). One of the most significantly downregulated genes in IgE PBs was spleen associated tyrosine kinase (SYK), which plays an essential role in BCR signal transduction (30) and is necessary for naïve B cell differentiation into plasma cells and for memory B cell survival (31). Taken together, this gene expression program shows that the IgE PB cell state is immature relative to other PBs with weakened activation, proliferation, and survival capacity. It also provides a potential transcriptomic mechanism for the hypothesized short-lived IgE PB phenotype described in mouse models of allergy (15, 32).

We found human IgE B cells belonging to the naïve / memory subset were deficient in immunoglobulin heavy chain membrane exon splicing compared to other common isotypes. Furthermore, membrane exon splicing was detected at low levels in non-IgE PBs, but not in IgE PBs (Fig. 2C-D). In fact, the absence of mIgE splicing rendered us unable to assess the relative utilization of the two splice variants of mIgE known to have distinct signaling characteristics (33, 34). The lack of mature mIgE transcripts could be explained by poor processing of pre-mRNA (35) and is consistent with low IgE surface protein we measured by FACS; indeed, mIgE surface protein levels on true IgE B cells did not exceed those of some non-IgE B cells presumably displaying surface IgE as a result of CD23-mediated capture (fig. S2B). These results suggest that the scarcity of circulating memory IgE B cells *in vivo* could result from impaired membrane IgE expression that compromises IgE B cell entry into the memory compartment and/or memory B cell survival. Murine studies support such a hypothesis, having shown IgE responses are reduced by removal or modification of mIgE domains, but augmented by the exchange of these domains for those of IgG1, thereby suggesting that poor mIgE expression and signaling acts to restrict IgE levels (36, 37).

### Characterization of peanut-specific IgE and IgG4 antibodies

Surprisingly, our clonal analysis produced one CF of cells belonging to multiple individuals (CF1, Fig. 1E), which contained three IgE PBs from individual PA12 and three IgE PBs from individual PA13. The antibodies produced by these six cells were highly similar (Fig. 3A, fig. S7A-B) as all utilized the IGHV3-30*18 and IGHJ6*02 heavy chain genes as well as the IGKV3-20*01 and IGKJ2*01 light chain genes, with pairwise CDR3 amino acid sequence identity ranging from 65% to 94% for the heavy chain and 70% to 100% for the light chain. These antibodies were also highly mutated and enriched in replacement mutations within the complementarity determining regions of both chains (fig. S7C). In fact, compared to all other class switched antibodies, these were amongst the most mutated: the heavy chains were in the 76th percentile or above for mutation frequency, while all of the light chains were in the 96th percentile or above (fig. S7D).

**Fig 3.**
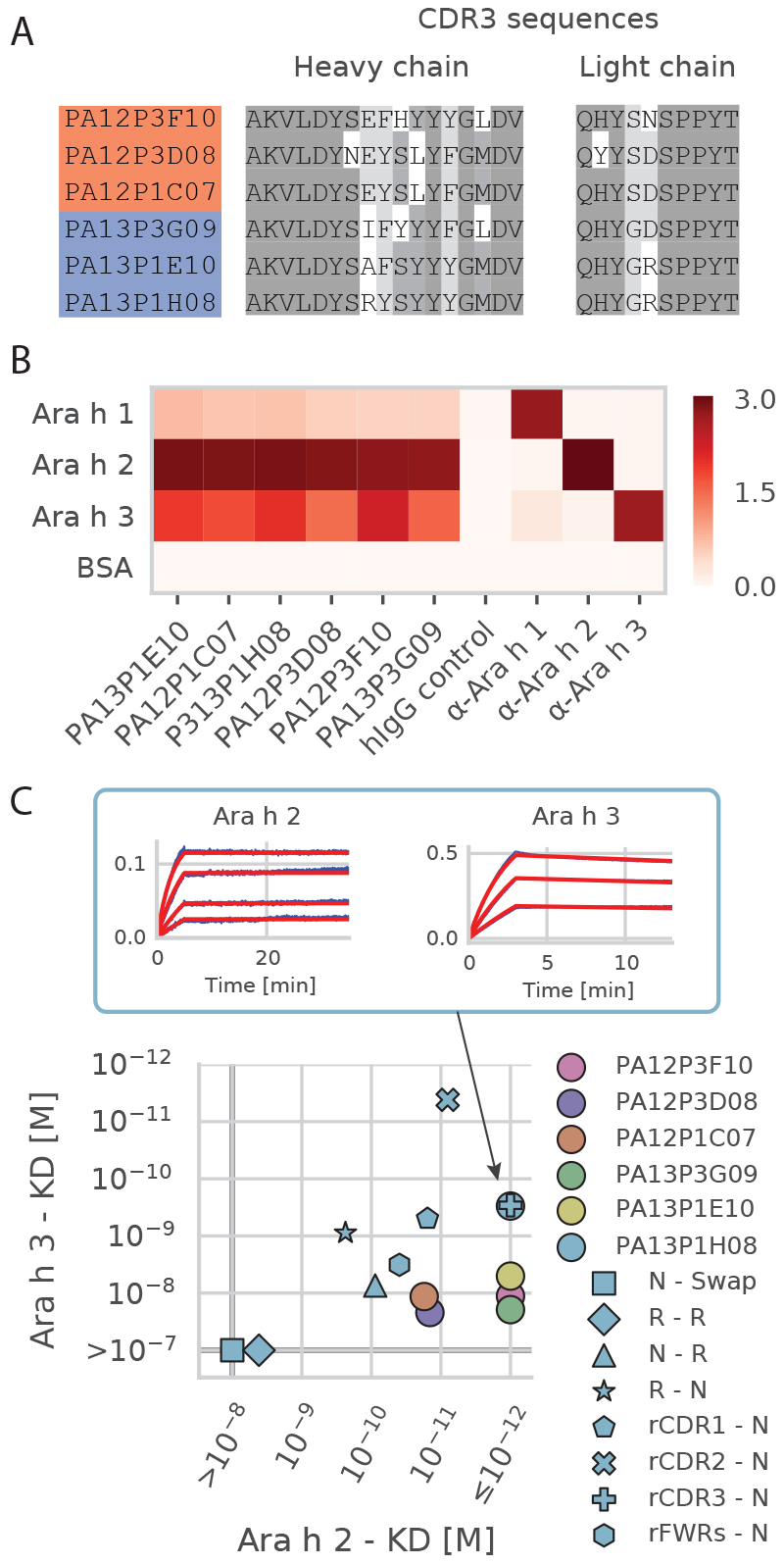
High affinity, cross-reactive human IgE antibodies. (A) Highly similar heavy and light chain CDR3s depict convergent evolution in two unrelated individuals (PA12 & PA13). Residues shaded by identity. (B) Indirect ELISA depicting antibody cross-reactivity to multiple peanut allergens. Commercially available mouse monoclonal αAra h antibodies served as positive controls. (C) Dissociation constants (KDs) to major allergenic peanut proteins Ara h 2 and Ara h 3 for each antibody as well as eight variants of PA13P1H08, designated as “heavy - light,” using the abbreviations: N=native, R=reverted, FWRs=framework regions. An “r” prefix indicates reversion of only that region. Binding curves for PA13P1H08 shown above.

We recombinantly expressed the six IgE antibodies belonging to this convergent clonal family in order to assess whether they bind the natural forms of the major allergenic peanut (*Arachis hypogaea*) proteins Ara h 1, Ara h 2, or Ara h 3. Of all characterized peanut allergens, Ara h 2 is the most commonly recognized by allergic individuals and is the most clinically relevant both in terms of immunological response (38) and discriminating allergic status (39, 40). Using an indirect ELISA as a qualitative screen for binding, we found that, surprisingly, all six antibodies were cross-reactive; they bound strongly to Ara h 2, moderately to Ara h 3, and very weakly to Ara h 1 (Fig. 3B). We then used biolayer interferometry to determine dissociation constants of each antibody to Ara h 2 and Ara h 3, with resulting affinities of tens of picomolar (pM) to sub-pM for Ara h 2 and tens of nanomolar (nM) to sub-nM for Ara h 3 (Fig. 3C, fig. S8); these affinities are comparable to some of the highest affinity native human antibodies discovered against pathogens such as HIV, influenza, and malaria (41–45). Furthermore, if the antibodies we discovered or variants thereof were to be used therapeutically as blocking antibodies intended to outcompete endogenous IgE for allergen, an approach recently shown to be efficacious for treatment of cat allergy (46), such high affinity to multiple peanut allergens should be advantageous.

To investigate the degree to which each chain and the mutations therein affect antibody binding properties, we recombinantly expressed eight variants of antibody PA13P1H08, each with one or more regions in the heavy and/or light chain reverted to the inferred naïve rearrangement (fig. S7E-G). Retaining the native heavy chain while swapping the light chain with another kappa light chain from an antibody without peanut allergen specificity abrogated binding to both allergenic proteins, while reverting both chains largely eliminated Ara h 3 binding and dramatically reduced Ara h 2 affinity (Fig. 3C, fig. S8C). Reverting only the heavy or light chain reduced the affinity to Ara h 2 and Ara h 3, but disproportionately; light chain mutations contributed more to Ara h 3 affinity than did heavy chain mutations. We also found a synergistic contribution of heavy chain mutations to affinity as independent reversion of the CDR1, CDR2, or framework regions each caused minor decreases in affinity. Reversion of the CDR3 did not alter binding, which was not surprising given the two amino acid difference between the native and reverted CDR3 sequences. Interestingly, reversion of the heavy chain CDR2 increased Ara h 3 affinity, while only marginally decreasing Ara h 2 affinity. Together, these results indicate that while the inferred naïve antibody is capable of binding the most clinically relevant peanut allergen Ara h 2, mutations in both heavy and light chains are necessary to produce the high affinity and cross-reactive antibodies that we found in circulating IgE PBs of unrelated individuals.

We also expressed antibodies from two other CFs. CF2 contained three IgE PBs from individual PA16 (two of which were identical), but these antibodies did not bind Ara h 1, 2, or 3, which was unsurprising given this individual had low plasma peanut-specific IgE levels as well as IgE specific to other allergens (fig. S1). On the other hand, CF3 contained an IgE PB (PA15P1D05) and IgG4 PB (PA15P1D12), the recombinantly expressed antibodies from which did not bind Ara h 1 appreciably, but bound Ara h 3 with nanomolar affinity and Ara h 2 with sub-nanomolar affinity (fig. S8). Interestingly, these two antibodies utilize the same light chain V gene and a highly similar heavy chain V gene (IGHV3-30-3*01) as the six convergent antibodies of CF1, which provides support for the importance of these V genes in Ara h 2 and Ara h 3 binding. Moreover, the presence of peanut-specific IgE and IgG4 PBs in the same CF within an allergic individual provides a unique example of *in vivo* competition for allergen between two antibody isotypes with possible antagonistic effector functions in allergic disease.

### Biallelic and polarized germline transcription in single cells

Tailored responses of the adaptive immune system are possible in part due to the ability of activation-induced cytidine deaminase (AID) to initiate class switch recombination (CSR) in B cells, leading to the production of antibodies with specific effector functions. CSR is preceded by cytokine-induced germline transcription, where nonproductive germline transcripts (GLTs) that contain an I-exon, switch (S) region, and heavy chain constant region exons guide AID to the S region (47). Importantly, GLT processing is necessary for CSR (48, 49) and canonically results in two species: an intronic S region lariat and a mature polyadenylated transcript consisting of the I exon spliced to the constant region exons (50). In our scRNA-seq data, we observe multiple splice isoforms of the latter, where the proximal constant region exon serves as the exclusive splice acceptor for multiple splice donors. IgE had the largest number of distinct GLTs at five (Fig. 4A and fig. S9), which we confirmed through Sanger sequencing (fig. S10); these were expressed in numerous cells of varying isotypes and across all individuals, but at nonuniform frequencies. The I-exon was the most common splice donor site (Fig. 4A, GLT #1) and it is known that I-exons can provide multiple splice donors (51–53), but ɛGLT splice donors within the switch region were also observed. We also found independent evidence for multiple ɛGLT splice donors in a previously published scRNA-seq dataset from murine B cells harvested 24 h after simulation to class switch (54) (fig. S11).

**Fig. 4.**
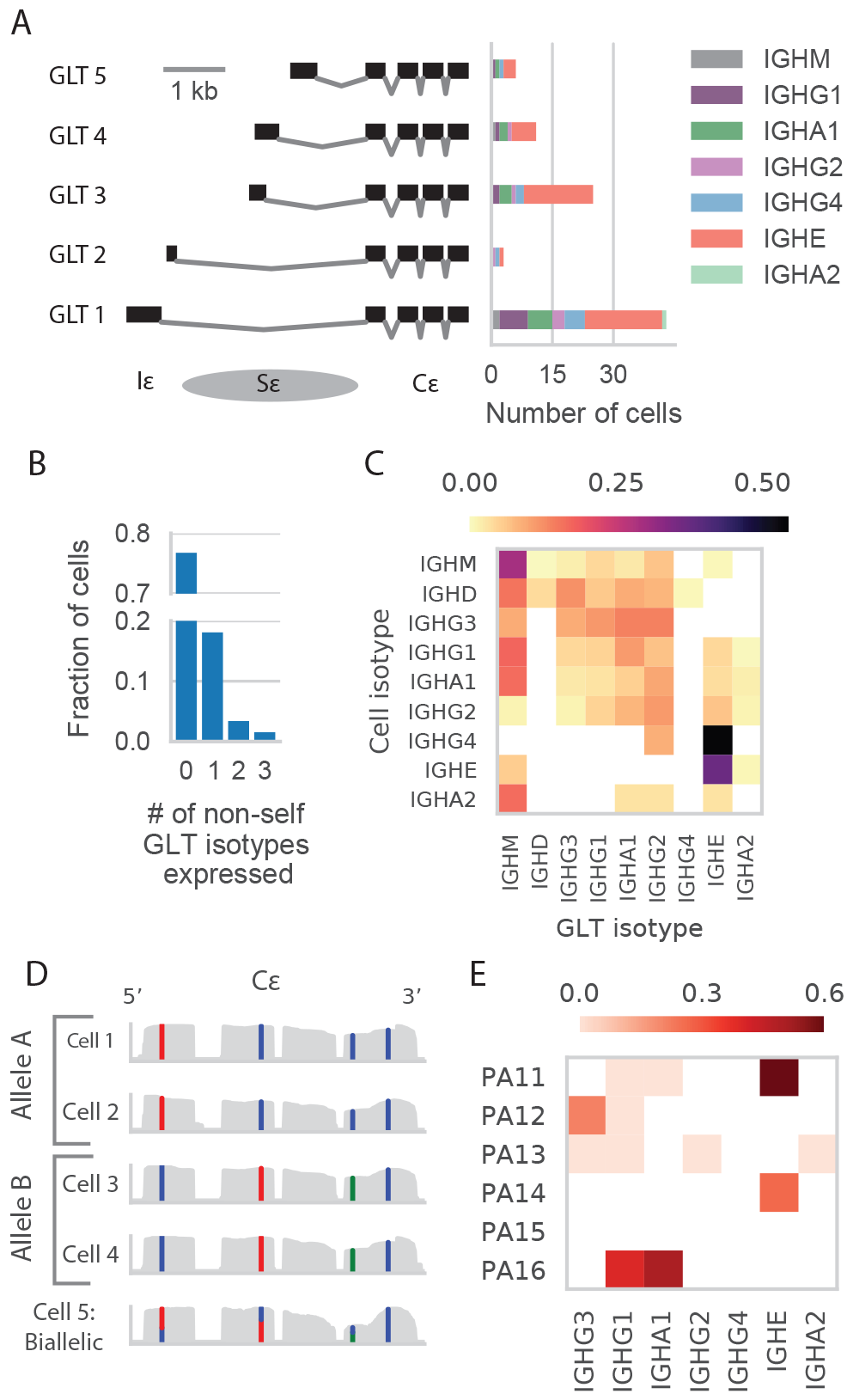
Germline transcription in single B cells. (A) Identification of Cε germline transcript splice donors along with the number of cells, by isotype, expressing each. (B) Histogram of the number of non-self GLT isotypes expressed in each cell. (C) Germline transcription heatmap indicating the fraction of cells of a given isotype (rows) expressing a given GLT (column). (D) Example from individual PA11 confirming biallelic ɛGLT expression in a non-IgE B cell after haplotype phasing the IgE constant region using single IgE B cells. (E) Heatmap indicating the fraction GLT-expressing cells for which expression is biallelic, by individual and GLT isotype. Analysis is limited to constant regions within individuals for which haplotype phasing could be performed.

We next assessed GLT expression across all isotypes. Most B cells did not express a GLT of a non-self isotype, while, unlike previous reports (55), those that did tended to be polarized towards the expression of a single GLT isotype (Fig. 4B). GLT production varied dramatically both by GLT isotype produced and by the cell’s current isotype (Fig. 4C). We observed a high proportion of IgG4 and IgE cells expressing ɛGLTs, while in contrast, we found almost no IgG4 GLT expression within any cells in these allergic individuals. Interestingly, we observed that GLT expression arising from the alternate allele is common, as evidenced by widespread expression of IgM GLTs in class switched B cells, and in some cases, expression of GLTs of isotypes upstream of a cell’s own class switched isotype (signal below the diagonal in Fig. 4C). Mirroring the landscape of human class switching (56), we observe the trend for GLT production to be higher for proximal downstream isotypes rather than distant downstream isotypes.

The study of B lymphocyte transcriptomes at single cell resolution offers other advantages; for example, we discovered multiple instances of biallelic GLT expression within single cells through heavy chain constant region haplotype phasing in individuals who had heterozygous single nucleotide variants within these loci. An example of this process that demonstrates biallelic ɛGLT expression is shown in Fig. 4D. Not all constant regions within individuals had such enabling variants, but in those that did we observed biallelic expression was relatively common relative to monoallelic expression (Fig. 4E).

## Conclusion

Using scRNA-seq, we provide the first transcriptomic characterization of circulating human IgE B cells and the antibodies they produce. Our data suggest two mechanisms underlying IgE regulation in humans: a relative deficiency of membrane immunoglobulin expression and an immature IgE PB gene expression program suggestive of weakened activation, proliferation, and survival capacity. These results are largely consistent with extensive studies of mIgE signaling and IgE memory in murine models of allergy (57–61), and provide evidence supporting the use of animal models for this disease. Furthermore, the ability to capture GLT splice variant, polarization, and biallelic expression information within single B cells presents an exciting application of scRNA-seq for future mechanistic studies of GLT production and CSR.

Insight into convergent evolution of high affinity antibodies in unrelated individuals can guide vaccine design and lead to strategies for population-level passive immunity; it is also a process that has been argued to occur in response to a number of pathogens such as influenza (62), HIV (45), *Streptococcus pneumoniae* (63). Here we found a striking case of convergence where two unrelated individuals produced high affinity, cross-reactive, peanut-specific antibodies comprised of identical gene rearrangements within respective heavy and light chains. A third individual had Ara h 2-specific antibodies that utilized a similar heavy V gene and the same light chain V gene. Although our study was limited by sample size, there is evidence supporting the importance of these genes within the peanut-allergic population more broadly: one independent study of IgE heavy chain sequences from peanut allergic individuals (64) reported IgE heavy chains that utilized identical V and J genes and shared at least 70% CDR3 identity with one or more of the six convergent antibodies in our dataset (fig. S12); another study (9) reported Ara h 2 specific IgG and IgM antibodies that utilized similar IGHV3-30 genes.

Cross-inhibition experiments with purified allergens and plasma IgE have shown that crossreactivity of IgE antibodies may also be common within peanut allergic individuals (65) and the antibodies we have isolated here offer a clear example of these findings. Furthermore, the fact that these high affinity antibodies were being produced by secretory IgE PBs found in circulation contributes to an understanding of how minute amounts of allergen are capable of eliciting severe allergic reactions. We also expect that either these antibodies or engineered variants of them could be used as therapeutic agents; recent clinical results have shown that engineered allergen-specific IgG antibodies can be administered to humans and provide effective treatment for cat allergies, perhaps by outcompeting the native IgE for antigen (46).

## Acknowledgements

We would like to acknowledge helpful discussions with Felix Horns. This research was supported by the Simons Foundation (SFLIFE #288992 to SRQ), the Chan Zuckerberg Biohub, and the Sean N Parker Center for Allergy and Asthma Research at Stanford University. DC is supported by an NSF Graduate Research Fellowship and the Kou-I Yeh Stanford Graduate Fellowship.

## Data availability

Processed data including antibody assembly information and the gene expression count matrix is available in the Supplementary Materials. Raw sequencing data is available from the Sequence Read Archive (SRA) under BioProject accession PRJNA472098.

## Material and Methods

### Study subjects

All study subjects were consented and screened through a Stanford IRB approved-protocol. Participants were eligible if they had a peanut allergy confirmed by an oral food challenge and board certified allergist. Peanut allergic individuals with reported reactivity to peanut ranged in age from 7 to 16, and in some cases exhibited sensitivities to other food allergens (fig. S1).

### Plasma IgE measurement and B cell isolation

Both plasma and cellular fractions were extracted from up to 45 mL of fresh peripheral blood collected in K_2_ EDTA tubes. For plasma extraction, blood was transferred to 15 mL falcon tubes and spun at 1600 g for 10 min. The upper plasma layer was extracted, transferred to 2 mL Eppendorf protein LoBind tubes and spun again at 16000 g to further purify the plasma fraction. The resulting supernatant was moved to fresh tubes before being put on dry ice and later transferred to −80°C. Allergen-specific plasma IgE measurements were performed by CLIA-licensed Johns Hopkins University Dermatology, Allergy, and Clinical Immunology (DACI) Reference Laboratory using the ImmunoCAP system. To purify B cells remaining after plasma extraction, RosetteSep human B cell enrichment cocktail (Stemcell Technologies), a negative selection antibody cocktail, was added after the plasma fraction was replaced with PBS + 2% fetal bovine serum (FBS). After a 20 min incubation, the blood was then diluted two-fold with

PBS + 2% FBS before being transferred to Sepmate 50 mL tubes (Stemcell Technologies) containing 15 mL Ficoll-Plaque PLUS (GE Healthcare Life Sciences). An enriched B cell population was achieved after a 10 min, 1200 g spin with the brake on and transferred a fresh tube. Residual red blood cells were then removed using ACK lysis buffer (ThermoFisher) and cells were washed with stain buffer (BD Biosciences). Cells were stained on ice with the following BioLegend antibodies according to the manufacturer’s instructions: PE anti-human IgE clone MHE-18, Brilliant Violent 421 anti-human CD19 clone HIB19, APC anti-human IgM clone MHM-88, and Alexa Fluor 488 anti-human IgG clone M1310G05. Cells were washed twice more prior to sorting.

### Flow cytometry and single cell sorting

Single cell sorts were performed on a FACSAria II Special Order Research Product (BD Biosciences) with a 5 laser configuration (355, 405, 488, 561, and 640 nm excitation). Fluorophore compensation was performed prior to each sort using OneComp eBeads (ThermoFisher), although minimal compensation was required due to the fluorophore panel and laser configuration. Equivalent laser power settings were used for each sort. Cells were sorted using “single cell” purity mode into chilled 96 well plates (Biorad HSP9641) containing 4 μL of lysis buffer (66) with ERCC synthetic RNA spike-in mix (ThermoFisher). Plates were spun and put on dry ice immediately before storage at −80°C.

### cDNA generation, library preparation, and sequencing

A modified version of the Smart-seq2 protocol (11) was used as previously described (66), but with 25 cycles of PCR amplification due to the low mRNA content of naïve and memory B cells. In total, 1165 cells were sequenced across 5 runs using 2 × 150 bp Illumina High Output kits on an Illumina NextSeq 500.

### Sequencing read alignment, gene expression, and splicing

Sequencing reads were aligned to the genome in order to determine gene expression and identify splice variants. To produce the gene expression counts table, reads were first aligned to the GRCh38 human genome using STAR v2.5.3a (12) run in 2-pass mode. Gene counts were then determined using htseq-count (67) run in intersection-nonempty mode. The GTF annotation file supplied to both STAR and htseq-count was the Ensembl 90 release manually cleaned of erroneous immunoglobulin transcripts e.g. those annotated as either a V gene or constant region but containing both V gene and constant region exons. During STAR genome generation an additional splice junction file was provided that included splicing between all combinations of heavy chain CH1 exons and IGHJ genes to improve read mapping across these junctions. Gene expression was normalized using log2 counts per million after removing counts belonging to ERCCs. Cells with fewer than 950 expressed genes were excluded prior to analysis (fig. S3B), as were putative basophils, identified by high FACS IgE, absent or poor quality antibody assemblies, and expression of histidine decarboxylase (HDC) and Charcot-Leyden crystal protein/Galectin-10 (CLC). Batch effects mostly affecting the naïve / memory B cell subset were noted between sorts by clustering using PCA on the 500 most variable genes; this gene set was enriched in genes known to be affected by sample processing such as FOS, FOSB, JUN, JUNB, JUND, HSPA8 (68). PCA following the exclusion of genes differentially expressed between sort batches (Mann-Whitney test, p-value < 0.01 after Bonferroni correction) yielded well-mixed populations within both the naïve / memory and PB cell clusters not biased by sort batch, individual, or sequencing library (fig. S3G). For differential expression analysis between IgE and non-IgE PBs, genes expressed in at least 10 PBs were analyzed by voom-limma (69) with sort batch and sequencing library were supplied as technical covariates. Constant region genes, such as IGHE and IGHA1, were excluded given these are differentially expressed by design of the comparison being made.

Analysis of splicing, including GLT expression, relied upon splice junctions called by STAR. Junctions were discarded if they contained fewer than three unique reads and GLT splice donors were only considered if observed in at least three cells. It should be noted the elevated levels of IgG2 GLT production can be explained by splicing of the CH1 IgG2 exon to an upstream lincRNA (ENSG00000253364). Biallelic expression of GLTs was determined based on heterozygous expression of single nucleotide variants discovered within heavy chain constant regions using bcftools (70). For the analysis of immunoglobulin heavy chain constant region exon splicing and coverage, genomic coordinates from the Ensembl gene annotation were used. Read coverage of these exons was generated using the samtools (71) depth command. To illustrate the absence of IgE membrane exon coverage, cells were leniently considered to have “any” membrane exon coverage (Fig. 2C-D) if at least 5% of either membrane exon had at least 5 reads.

### Antibody heavy and light chain assembly

In addition to alignment, sequencing reads were also independently assembled in order to reconstruct full length heavy and light chain transcripts. BASIC (72) was used as the primary assembler given its intended use for antibody reconstruction, while Bridger (73), a *de novo* whole transcriptome assembler, was used to recover the minority of heavy and/or light chains lacking BASIC assemblies. The heavy chain isotype or light chain type (lambda or kappa) was determined using a BLAST (74) database of heavy and light chain constant regions constructed from IMGT sequences (75). Immunoglobulin variable domain gene segment assignment was performed using IgBLAST (76) v1.8.0 using a database of human germline gene segments from IMGT. IgBLAST output was parsed with Change-O and mutation frequency was called with SHazaM (77). Cells without a single productive heavy and single productive light chain, which were all members of the naïve / memory cell cluster, were excluded, leaving a final total of 973 cells. Graph-tool (https://graph-tool.skewed.de/) was used to draw clonal families and the workflow engine Snakemake (78) was used to execute all analysis pipelines.

### Recombinant antibody expression

Recombinant expression of select antibodies enabled characterization of antibody specificity and affinity. All heavy chains were expressed as human IgG1, while light chains were expressed as either lambda or kappa as appropriate. Heavy and light chain sequences were synthesized by Genscript after codon optimization and were transiently transfected in HEK293-6E cells. Antibodies were purified with RoboColumn Eshmuno^®^ A columns (EMD Millipore) and were confirmed under reducing and non-reducing conditions by SDS-PAGE and by western blots with goat anti-human IgG-HRP and goat anti-human kappa-HRP or goat anti-human lambda-HRP as appropriate.

### Functional antibody characterization

ELISAs were performed to qualitatively assess peanut allergen binding. Purified natural Ara h 1 (NA-AH1-1), Ara h 2 (NA-AH2-1) and Ara h 3 (NA-AH3-1), purchased from Indoor Biotechnologies, were immobilized overnight at 4°C using 50 μL at a concentration of 2 ng / uL. Following 3 washes, wells were blocked with 100 μL of PBST (ThermoFisher) + 2% BSA for 2 hours. After two washes, 100 μL of primary antibodies were incubated for 2 hours at a concentration of 2 ng / μL in blocking buffer. Following 4 washes, 100 μL of rabbit anti-human HRP (abcam #ab6759) or rabbit anti-mouse HRP (abcam #ab6728) secondary antibodies were incubated for 2 hours at a dilution of 1/1000 in blocking buffer. After 5 washes, 150 μL of 1-Step ABTS Substrate Solution (ThermoFisher) was added to the wells. Color development was measured at 405 nm on a plate reader after 8 - 20 min and reported OD values are after subtraction of signal from no-antibody wells. Negative controls included immobilized BSA as an antigen, as well as a human isotype control primary antibody (abcam #ab206195). One random IgM / IgK antibody we expressed (PA12P4H03) also did not exhibit any binding. Positive controls consisted of monoclonal mouse antibodies 2C12, 1C4, and 1E8 (Indoor Biotechnologies) specific for Ara h 1, Ara h 2, and Ara h 3, respectively.

Kinetic characterization of antibody interactions with natural purified allergenic peanut proteins was achieved using biolayer interferometry on a ForteBio Octet 96 using anti-human IgG Fc capture (AHC) biosensors with 1X PBST as the assay buffer. The assay was run with the following protocol: up to 600s baseline, 120-150s antibody load, 120-300s baseline, associations of up to 300s, and variable length dissociations that lasted up to 30 min for high affinity antibody-antigen interactions. Biosensors were regenerated by cycling between buffer and pH 1.5 glycine following each experiment. Antibodies were loaded at a concentration of 10 nM, while optimal peanut protein concentrations were determined experimentally (fig. S8). Data were processed using ForteBio software using a 1:1 binding model and global fit after reference sensor (ligand, but no analyte) subtraction

## Supplementary figure captions

**Fig. S1. Plasma IgE levels.** (A) Allergen-specific and allergen component (hazelnut, peanut) concentrations. (B) Total IgE concentration. (C) Positive correlation between total plasma IgE concentration and the frequency of IgE B cells among CD19+ B cells. Each point is an individual.

**Fig. S2. FACS gating and analysis.** (A) Gating strategy for sorting single B cells. IgE+ B cells have been overlaid as red dots. (B) Isotype identity within the final IgE gate as determined by heavy chain transcript assembly. ND=not determined. (C) For reference, putative basophils (CD19- IgE+) display higher IgE surface expression than IgE+ B cells.

**Fig. S3. scRNA-seq data overview and quality control.** (A) Cells were sequenced in 5 libraries to a depth of ~1-2 million reads / cell. (B) Genes per cell histogram. Cells expressing fewer than 950 genes were discarded. (C) Rarefaction curve depicting the number of genes detected as a function of sequencing depth for eight randomly selected cells in each B cell subtype. Solid lines and shaded area represent mean and 95% confidence interval for the gene count, respectively. (D) Read mapping distribution for retained cells. Most reads mapped uniquely (Ensembl gene annotation) and multimapped reads largely belonged to RNA18S5 repeats on chr21 and unplaced scaffolds. (E) Read mapping across gene bodies showed minimal 3’ or 5’ bias. (F) V gene assembly length histogram by chain. (G) PCA on the top 500 most variable genes before (top) and after (bottom) batch correction.

**Fig S4. Auxiliary data supporting B cell subtype classification.** PBs (blue) have greater FACS forward and side scatter (A), more cDNA after Smart-seq2 preamplification (B), and have greater gene expression of antibody light and heavy chain constant regions (C) as compared to the naïve / memory B cell subset (pink). (D) Top differentially expressed genes for each subset.

**Fig. S5. Additional single cell characterization and analysis of clonal families.** (A) PCA plot as in Fig. 1B, but colored by isotype. (B-D) Analysis of clonal families (CFs). (B) Distribution of the number of cells per CF. (C) Fraction of cells of each isotype that belong to a multimember CF. (D) Heavy (right, blue) and light (left, red) chain CDR3 sequences and similarity heatmap for CFs in Fig 1E.

**Fig S6. Antibody comparisons within the IgE isotype and across isotypes**. (A) Heatmap indicating number of IgE antibodies with a given heavy and light chain CDR3 length. (B) Heavy and light chain mutation frequency of each IgE antibody. (C) Silent (S) and replacement (R) mutations by region within IgE heavy and light chains. (D) Heavy chain mutation frequency by isotype. (E) Relative utilization of the lambda and kappa light chain by isotype. (F) Heavy chain V and J gene usage heatmaps by isotype.

**Fig S7. Sequences, characteristics, and engineered variants of convergent IgE antibodies belonging to CF1.** (A-D) Antibody colors are conserved among panels. (A) Heavy chain amino acid sequences and the inferred naïve rearrangement (“Reverted”). Residues shaded by identity. (B) As in (A), but for the light chain. (C) Frequency of silent (S) and replacement (R) mutations by region. (D) Mutation frequency percentiles compared to all class-switched antibodies. (E-G) Engineered variants of PA13P1H08. (E) Native and reverted heavy chain sequences, in addition to sequences where region(s) of the heavy chain have been reverted to the inferred naïve rearrangement. Labels with an “r” prefix indicate only that region has been reverted. FWRs = frameworks. (F) Native and reverted light chain sequences. (G) Sequence of a light chain taken from a random antibody, PA12P4H03, which did not bind any peanut allergens by ELISA.

**Fig S8. Antibody specificity and affinity measurements.** (A) Ara h 2 binding curves acquired using biolayer interferometry. Each plot depicts a serial two-fold dilution starting with the allergen concentration inscribed in the upper right. (B) As in (A), but for Ara h 3. (C) Indirect ELISA of PA13P1H08 variants. For reference, names are designated as “heavy - light,” using the abbreviations: N=native, R=reverted, FWRs=framework regions. An “r” prefix indicates reversion of only that region.

**Fig S9. GLT splice donors for all isotypes.** Note that only the first three constant region exons of each isotype are shown for clarity.

**Fig. S10. Sanger sequencing of five ɛGLTs amplified from single cell cDNA confirms GLT identity and splicing.** Shown for each is of 70 nt of GLT sequence spliced to the first 70 nt of IGHE CH1 in the 5’ ↓ 3’ orientation. For each GLT, the expected upper sequence agrees with the lower Sanger sequencing result. The same reverse IGHE CH2 primer (5’ - TTGATAGTCCCTGGGGTGTACC - 3′) was used for all GLTs along with the following forward primers (5’ –> 3’): (A) CTGGACTGGGCTGAGCTAGAC, (B-C) GGCCTGAGCTGTGATTGGAAG, (D) CACCCTCACAGCATCAACCAAG, (E) TGCCCGGCACAGAAATAACAAC.

**Fig S11. Stimulated murine B cells produce multiple ɛGLTs.** IGV coverage histograms and splice junctions for the murine ighe constant region in single cells stimulated with IL-4, LPS, and BAFF (54). Arrows indicate unique ɛGLT splice donors.

**Fig S12. Similar IgE heavy chain CDR3 sequences in an independent dataset**. Pairwise CDR3 sequence identity of the six convergent heavy chain CDR3 sequences from CF1 of the present study and three IgE heavy chain CDR3 sequences derived from multiple patients in a separate peanut allergy immune repertoire sequencing study (64). Each CDR3 sequence from this separate study shares at least 70% identity with one or more CDR3 sequences from the present study. All sequences share the IGHV3-30 and IGHJ6 gene segments and have CDR3s 17 amino acids in length.

Supplementary tables

**Table S1. Gene expression count matrix.** Each column is a cell and each row is a gene.

**Table S2. Single cell antibody assemblies.** V, (D), and J gene segment calls as well as isotype and CDR3 amino acid sequence for the heavy and light chains of each cell.

